# Inhibition of CD40-TRAF6 signaling protects against aneurysm development and progression

**DOI:** 10.1101/2023.03.24.534110

**Authors:** Miriam Ommer-Bläsius, Tanja Vajen, Christin Elster, Sarah Verheyen, Susanne Pfeiler, Christine Quast, Julia Odendahl, Alexander Lang, Malte Kelm, Esther Lutgens, Norbert Gerdes

## Abstract

**Objective:** Inflammation is a critical process during the progressive development and complication of abdominal aortic aneurysm. The co-stimulatory dyad CD40-CD40L is a major driver of inflammation and modulates immune responses. This study evaluates the potential of a small molecule inhibitor, which blocks the interaction between CD40 and tumor necrosis factor (TNF) receptor-associated factor (TRAF)-6, referred to as TRAF-STOP, in the early and later phase during AAA progression.

**Methods and results:** AAAs were induced in C57BL/6J mice by infrarenal aortic porcine pancreatic elastase infusion for 7, 14 or 28 days. Inhibition of CD40 signaling by TRAF-STOP resulted in less severe AAA formation and reduced the incidence of AAA development. TRAF-STOP treatment attenuated aortic structural remodeling, characterized by a reduced elastic fiber degradation, lowered expression of matrix metalloproteinase (MMP)-2 and MMP9, as well as preserved collagen type IV content in aneurysmal tissue. Furthermore, this is accompanied by the reduction of key pro-inflammatory genes such as TNFα.

**Conclusion:** Pharmacological inhibition of CD40-TRAF6 signaling protects from adverse aortic structural remodeling during the early phase of AAA progression representing a translational strategy to limit progression of human AAA disease.

## 1. Introduction

Abdominal aortic aneurysm (AAA) is a life-threatening disease, defined by a persistent dilatation of the abdominal aorta by >3cm or >50%. Especially men over the age of 65 years show a high prevalence of up to 8% compared to a heterogeneous range of 0.37% to 1.53% in women over the age of 60 years.^1, 2^ In the event of an aortic rupture or dissection, AAA is associated with a high acute-mortality rate of >80%.^3^ Although major advances in surgical and endovascular repair of AAA has been achieved, there is no drug-based therapy available.^4^ The main pathological hallmarks of AAA include extracellular matrix (ECM) remodeling that is associated with degeneration and loss of vascular smooth muscle cells and the accumulation and activation of inflammatory cells. The inflammatory process is of major importance in the progression and development of AAA, as it essentially influences many determinants of aortic wall remodeling. The infiltration and activity of both innate and adaptive immune cells contributes to aortic aneurysm development.^5^

Patients with AAA show increased serum concentrations, as well as an upregulated expression in aneurysmal tissue of pro-inflammatory cytokines such as interleukin-1 (IL-1)-β, IL-6, tumor necrosis factor TNF)-α, and interferon (IFN)-γ.^3, 6^ In mice, pro- and anti-inflammatory mediators such as IFNγ, TNFα and transforming growth factor (TGF)-β, respectively affect the development of AAA.^7, 8^ Elastin and collagen are the primary extracellular matrix components of the aortic wall. Elastic fibers provide reversible elasticity to the aorta, while collagen fibers provide strength and limit distension at high pressures.^9^ During AAA development expression and activity of matrix metalloproteinases (MMP) such as MMP2 and MMP9 increase, promoting extracellular matrix and elastin degradation, thus provoking aortic dilation. Furthermore, deficiency in these enzymes protect from AAA formation.^10-13^ The co-stimulatory dyad CD40-CD40L contributes to inflammatory and auto-immune-related diseases such as atherosclerosis, Crohn’s disease, encephalitis and arthritis.^14^ In previous studies, we identified a central role for CD40L in the development and complications of AAA. CD40L deficient mice were protected from dissecting aneurysm formation and showed a decrease in the incidence of fatal rupture due to a decreased accumulation and activation of inflammatory cells and a reduced MMP2 and MMP9 activity in the arterial wall.^15^ Moreover, interfering with CD40 signaling in an experimental atherosclerosis model could attenuate plaque progression by reducing leukocyte infiltration and cytokine release.^16^

In this study, we investigate the pharmacological inhibition of CD40 signaling via its adaptor molecule TNF receptor associated protein (TRAF)-6, that is predominantly utilized in monocytes and macrophages. Using the small molecule inhibitor TRAF-STOP we assess whether and how interruption of this specific pathway affects the development and progression of AAA formation in mice.

## 2. Materials and methods

### 2.1 Mice

Male C57BL/6J mice were purchased from Janvier Labs (Saint-Berthevin Cedex; France). All mice were about 10 weeks of age at the beginning of the experiments and received standard chow and drinking water *ad libitum*. Mice were kept in climatized cages with a 12-h–light/dark cycle. All animal experiments were performed according to ARRIVE (Animal Research: Reporting of In Vivo Experiments) II guidelines and approved by LANUV (North Rhine-Westphalia State Agency for Nature, Environment and Consumer Protection, file number AZ 81-02.04.2018.A408) in accordance with the European Convention for the Protection of Vertebrate Animals used for Experimental and other Scientific Purposes.

### 2.2 Experimental aneurysm induction and pharmacological inhibitor treatment

In this study, AAA formation was induced by aortic perfusion with porcine pancreatic elastase (PPE). Briefly, an isolated infrarenal aortic segment of C57BL/6J mice was perfused with sterile isotonic saline containing type I PPE (2-3 U/ml, E1250, Sigma-Aldrich, Burlington, MA, USA) for 5 min under pressure (120 mmHg) [12]. Depending on the batch, elastase concentrations vary from 2-3 U/ml.

In this study CD40-TRAF6 signaling was inhibited using the small molecule TRAF-STOP 6860766, further referred to as TRAF-STOP, previously described by Seijkens et al. [16]. On the day of AAA induction, mice were injected intraperitoneally (i.p.) with either TRAF-STOP at a final concentration of 10 µmol/kg/injection or 2.5% dimethyl sulfoxide (DMSO, ThermoFisher Scientific, Waltham, USA) in PBS as solvent control. Treatment was carried out three times per week. At the end of the experimental period (d7, d14 or d28), mice were euthanized with Ketamin (100 mg/kg; Ketaset™, Zoetis, USA) and Xylazin (10 mg/kg; Rompun™, Bayer, Germany).

### 2.3 Ultrasound imaging

AAA progression was measured weekly using non-invasive ultrasound technology. The Vevo 3100 high-resolution *in vivo* imaging system with a 25-55 MHz - transducer (MX550D) (VisualSonics Inc., FUJIFILM, Toronto, Canada) was used to take images of the aorta. For imaging mice were anesthetized with isoflurane and placed on a heated pad at 37°C. During the whole set of measurements body temperature, aspiration rate and electrocardiogram (ECG) were continuously monitored. Longitudinal B mode images of the infrarenal abdominal aorta were captured to assess aortic diameter. Aortic diameter was analyzed from leading to leading edge (LTL) in three cardiac cycles at end-diastole using Vevo LAB 5.6.0 software.

### 2.4 Aorta embedding and cutting

Aortae were harvested, fixed in 4% paraformaldehyde (PFA, J61899.AP; ThermoFisher Scientific) solution for 1 h at room temperature and dehydrated in 30% sucrose (S1888; Sigma-Aldrich) solution at 4°C. Thereafter, aortae were embedded in Tissue-Tek optimum cutting temperature (O.C.T.) freezing compound (Sakura Finetek, Amsterdam, The Netherlands). Cross-sections of aortic tissue were prepared with 5-7 µm thickness.

### 2.5 Histological staining

The cross-sections of aortic tissue at its largest diameter and three adjacent sections from each side were used for Elastika van Gieson staining. Briefly, staining of elastic fibers was performed by incubating the sections in Resorcinol-Fuchsine-solution (X877.1, Carl Roth, Karlsruhe, Germany) for 10 min, followed by a washing step under tap water for 30 sec. Afterwards, sections were stained with hematoxylin solution according to Gill II (T864.2, Carl Roth) for 30 sec, followed by a blueing step under running water for 1 min. Thereafter, the sections were stained with Picofuchsin using Van Gieson’s solution kit (3925.1, Carl Roth) for 2 min. After washing the sections with deionised water, they were mounted with VectaMount AQ Aqueous (H-5501, Vector Laboratories, Newark, USA). This staining was used to assess the AAA size, the grade of elastic fiber degradation and the aortic wall thickness. The elastin degradation was analyzed using a grading system with four grades, grade 1: completely intact fibers to isolated elastic fiber breaks; grade 2: increased amount of elastic fiber breaks to area-wide absence (max 30%) of 1 to 2 elastic fibers; grade 3: area-wide absence (30-50%) of 2 to 4 elastic fibers; grade 4: area-wide absence (more than 50%) of 2 to 4 elastic fibers or complete destruction of fibers. Overall elastic fiber degradation for one AAA was determined by calculating the average value of the 7 stained and graded cross-sections from one animal. AAA size and aortic wall thickness were analyzed using Fiji/ImageJ (V1.53t).

### 2.6 Immunofluorescence staining

For immunofluorescence staining of collagen type IV, 5-7 µm thick cross-sections of aortic tissue at its largest diameter and one adjacent section from each side were used. Unspecific binding sites were blocked by incubating the slides in DPBS supplemented with 0.1% Saponin Quillaja sp. (S4521; Sigma-Aldrich), 0.5% Bovine Serum Albumin (BSA; fraction V; 8076.3; Carl Roth) and 0.2% fish gelatine (G7765; Sigma-Aldrich) at room temperature for 1 h. Afterwards, cross-sections were stained with collagen type IV antibody (5µg/ml, polyclonal, ab6586; abcam, Cambridge, United Kingdom) overnight at 4°C. On the next day slides were incubated with a fluorescence labeled secondary antibody (Goat anti-Rat IgG (H+L) Cross-Adsorbed Secondary Antibody, Alexa Fluor™ 680, 2 µg/ml; A-21096, ThermoFisher Scientific) at room temperature for 1 h. Autofluorescence of aortic tissue was quenched using the Vector TrueVIEW Autofluorescence Quenching Kit (VEC-SP-8400, Vector Laboratories, Newark, USA). Slides were incubated for 5 min at room temperature. Nuclei were stained with 4, 6 diamidino-2-phenylindole dihydrochloride (DAPI, 1 ng/ml) for 5 min at room temperature. Finally, aortic sections were mounted with VectaMount AQ Aqueous. The analysis of the collagen amount in aortic tissue was performed semi-automated using a Fiji/ImageJ Macro (V1.53t). The average value of the three stained cross-sections were compared between treatment groups.

### 2.7 RNAscope technology

RNAscope™ *In situ* hybridization technique was used for spatial detection of RNA expression on 5-7 µm thick cryo-sections. Staining was performed with RNAscope™ target probes to Mm-Mmp2 (315931-C3), Mm-Mmp9 (472401-C2) Mm-TNFa (311081) on AAA tissue according to manufacturer’s instructions in the RNAscope™ Multiplex Fluorescent Detection Kit v2 (323110, ACD, Bio-Techne, Hayward, USA) using protease IV and a extended hybridisation time of 3h. RNA expression was quantified with the analyzing tool from LasX (Leica Microsystems, Wetzlar, Germany).

### 2.8 Microscopy

The fluorescence microscope DM6 B with a DFC9000 camera (Leica Microsystems) was used to capture images of stained aortic cross-sections.

### 2.9 Spatial transcriptomics

Aortic cross-sections were produced according to the instructions in section 2.4 Aorta embedding and cutting. The 5 µm slices were transferred onto Visium slides and stained with Hematoxylin and Eosin (H&E) according to procedure CG000160 Rev C.

Prepared Visium slides were used as input for the Visium Spatial Gene Expression workflow (10X Genomics, Pleasanton, USA) according to the manufacturer’s instructions. Aortic tissue permeabilization was carried out for 12 min. Sequencing was performed on a NextSeq 2000 system (Illumina Inc.; San Diego, CA, USA) with a mean sequencing depth of approximately 120.000 reads/spot. Raw sequencing data was processed using the 10X Genomics spaceranger software (v2.0.0). Alignment of reads to the mm10 genome, UMI counting as well as tissue detection and fiducial detection was performed via the *spaceranger* count pipeline. All samples were combined and normalized for sequencing depth through the spaceranger *aggr* pipeline.

### 2.10 Digestion of aortic tissue into single cells

AAA tissue was digested using an enzyme mixture adapted from^17^. Briefly, harvested aortae were dissected and digested in an solution containing 400 U/ml Collagenase I (C0130, Sigma-Aldrich), 120 U/ml Collagenase XI (C7657, Sigma-Aldrich), 60 U/ml Hyaluronidase I-S (H3506, Sigma-Aldrich) and 60 U/ml Dnase I (11284932001, Sigma-Aldrich) in DPBS containing calcium and magnesium (DPBS++, D8662, Sigma-Aldrich) supplemented with 20 mM HEPES (15630056, ThermoFisher Scientific) for 50 min at 37°C on a shaker at 600 rpm. Thereafter, cell suspension was poured onto a 100 µm cell strainer (130-110-917, Miltenyi Biotec, Bergisch Gladbach, Germany) and centrifuged for 10 min at 4°C and 450 x g.

### 2.11 Staining and sorting of single cells

Isolated aortic single cells were stained with Fc receptor blocker (TruStain FcX anti-mouse CD16/32; 101319; BioLegend, San Diego, CA, USA, 1:100) for 10 min at room temperature. Thereafter, cells were stained with TotalSeq™-A Mouse Universal Cocktail, V1.0 (dilution 1:1200; 199901; BioLegend) and a Hashtag antibody (TotalSeq™-B0303 and TotalSeq™-B0304) for 20 min at room temperature. Cells were centrifuged (5 min, 500 x g, 4 °C) and the supernatant was discarded. Afterwards single cells were resuspended in MACS buffer (130-091-221; Miltenyi Biotec) and stained with DAPI (1 ng/ml). Directly after adding DAPI to the cells, cells were sorted using a MoFlo XDP (Beckman Coulter, Krefeld, NRW, Germany). Viable CD45+ cells were collected and further used for Cellular Indexing of Transcriptomes and Epitopes by sequencing (CITESeq).

### 2.12 Single Cell Library Generation

Single Cell libraries were generated on the 10X Chromium Controller system utilizing the Chromium Next GEM Single Cell 5’ Kit v2 (10X Genomics) according to manufacturer’s instructions. Sequencing was carried out on a NextSeq 550 system (Illumina Inc.) with a mean sequencing depth of ∼50,000 reads/cell for gene expression, ∼20.000 reads/cell for the TotalSeq-ADT library and ∼5,000 reads/cell for the HashTag library.

For the TotalSeq-ADT library TotalSeq™-A Mouse Universal Cocktail, V1.0 was used according to manufacturer’s instructions. For the HashTag library TotalSeq™-B0303 anti-mouse Hashtag 3 Antibody (Clone M1/42; 30-F11; 155835; BioLegend) was used to mark single cells of DMSO treated mice and TotalSeq™-B0304 anti-mouse Hashtag 4 Antibody (Clone M1/42; 30-F11; 155837; BioLegend) was used for aortic cells of TRAF-STOP treated mice. Bioinformatica demultiplexing was performed with Seurat (v4.02) with the protocol “Demultiplexing with hashtag oligos” v. 2022-01-11.

### 2.13 Processing of 10X Genomics single cell data

Raw sequencing data was processed using the 10X Genomics CellRanger software (v6.0.2). Raw BCL-files were demultiplexed and processed to Fastq-files using the CellRanger *mkfastq* pipeline. Alignment of reads to the mm10 genome and UMI counting was performed via the CellRanger *multi* pipeline to generate a gene-barcode matrix.

### 2.14 Statistics

Data are presented as mean ± standard deviation (SD). P < 0.05 was considered as statistically significant. Kolmogorov-Smirnov test was used to test for normal distribution. Differences between two groups were evaluated using two-way repeated measures analysis of variance (ANOVA) with Sidak’s post test if data were normally distributed. Fisher’s exact test was used to determine the significant association between two categorical variables.

## 3 Results

### Inhibition of CD40 signaling by TRAF-STOP protects from elastase-induced AAA formation

To evaluate the potential of TRAF-STOP, a small molecule inhibitor that specifically blocks the interaction between CD40 and TRAF6, mice received solvent control or TRAF-STOP (10µmol/kg/injection) for 3 times per week starting on day 0 until day 7, 14 or day 28, respectively (**Figure 1A**). Following PPE infusion, the aortic diameter increases progressively from day 7 onward, and TRAF-STOP treatment significantly reduces the aortic dilation 14 and 28 days post AAA induction (**Figure 1B)**. Aneurysm formation and the change in aortic diameter was monitored and analyzed via ultrasound imaging (**Figure 1C**). An AAA was defined as an increase in AA diameter of at least 1.5 fold. Moreover, aortic cross-sections were histologically analyzed by Elastica van Gieson (EvG) staining to determine AAA size. Mice treated with TRAF-STOP developed overall smaller AAAs compared to DMSO-treated mice (**Figure 1D**). In the control group, 14 out of 16 mice (87.5 %) developed an AAA. In contrast, a reduced AAA incidence was observed in mice treated with TRAF-STOP (66.67 %) 28 days post AAA induction (**Figure 1E**).

**Figure 1.**
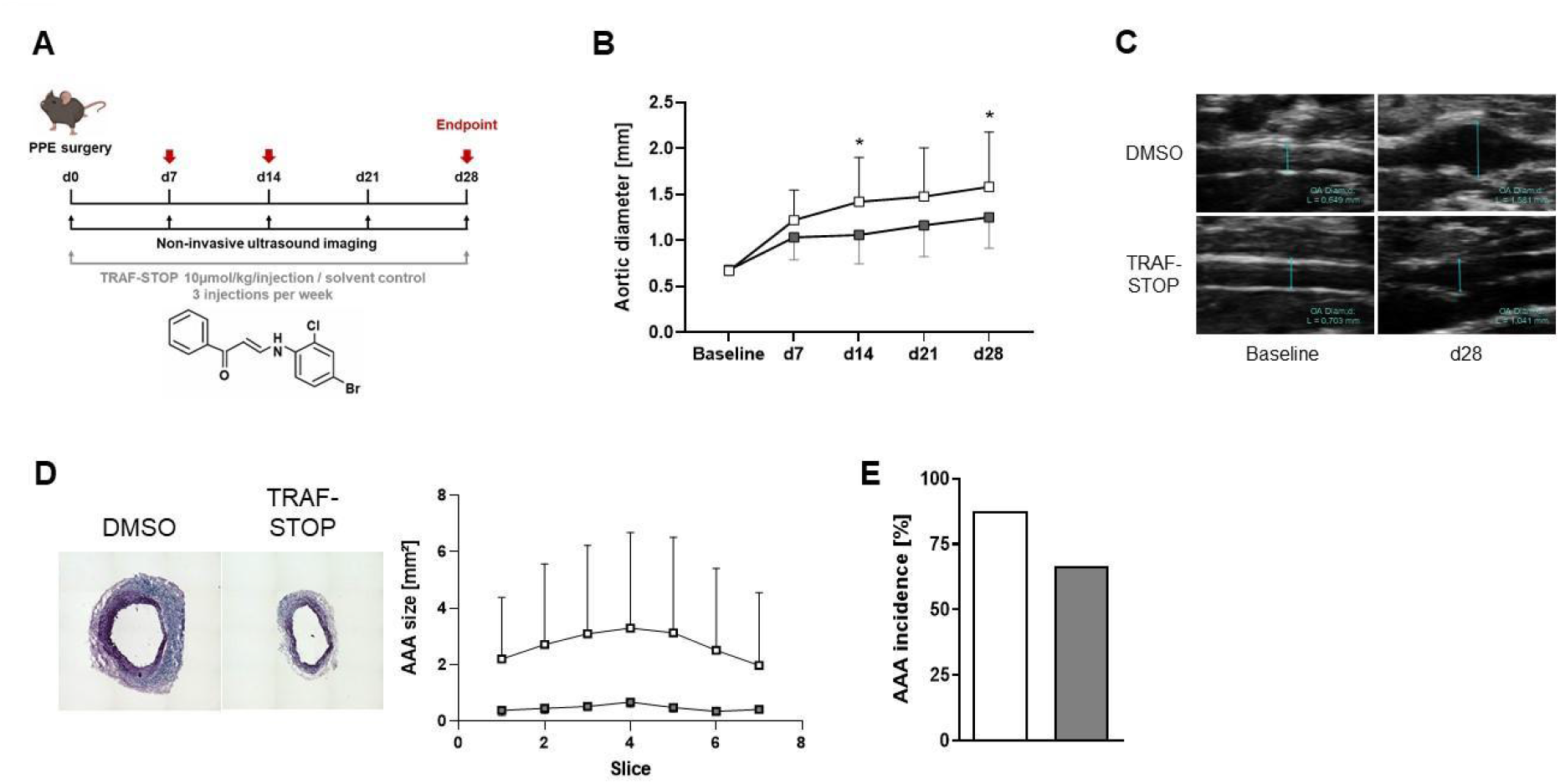
Inhibition of CD40 signaling by TRAF-STOP protects from elastase-induced AAA formation. (**A**) Experimental set-up and chemical structure of small molecule inhibitor, TRAF-STOP. (**B**) Abdominal aortic diameter was analyzed using ultrasound images. Dilation of abdominal aorta was significantly reduced in TRAF-STOP treated mice 14 and 28 days post PPE-induced AAA formation. 2-way ANOVA, Sidaks post test; *p<0.05; DMSO n=16, TRAF-STOP n=18. (**C**) Representative ultrasound images of the infrarenal aorta from mice treated with TRAF-STOP and solvent control prior to surgery and 28 days after AAA induction. (**D**) Representative Elastica van Gieson staining of elastic fibers in aneurysmal tissue 28 days post PPE-induced AAA formation and analyzed AAA size; DMSO n=4, TRAF-STOP n=4. (**E**) Decreased AAA incidence upon TRAF-STOP treatment 28 days after AAA induction. AAA incidence is defined as an increase in AA diameter of at least 1.5 fold. Two-sided Fisher’s exact test; p = 0.2327. DMSO n=16, TRAF-STOP n=18.

### Inhibition of CD40 signaling by TRAF-STOP protects from aortic structural remodeling by reduced of MMP2 and MMP9 expression and enhanced collagen degradation

EvG-stained cross-sections were analyzed to determine elastic fiber degradation and aortic wall thickness 28 days post PPE-induced AAA formation as an indication of disease progression. Mice treated with TRAF-STOP showed a decreased degradation of elastic fibers compared to DMSO-treated mice (**Figure 2A**). The degradation of elastic fibers was determined by a grading system of the elastic fiber destruction. In addition, TRAF-STOP-treated mice showed a decreased aortic wall thickness compared to DMSO treated mice (**Figure 2B**). Moreover, we investigated the content of collagen type IV in aneurysmal tissue by immunohistochemical staining, as collagen fibers are critical for aortic strength and stability. The collagen type IV amount in AAA cross-sections was increased 28 days after AAA induction upon TRAF-STOP treatment (**Figure 2C**). As MMP2 and MMP9 promote collagen and elastin degradation and thereby provoke aortic dilation, we investigated the mRNA expression of MMP2 and MMP9 by spatial transcriptomics 14 days after AAA induction. TRAF-STOP treatment reduced expression of MMP2 as well as MMP9 in aneurysmal tissue (**Figure 2D, E**). Furthermore, we validated these findings by performing RNAscope™ in AAA 14 days post PPE-induced AAA formation (**Figure 2D, E**).

**Figure 2.**
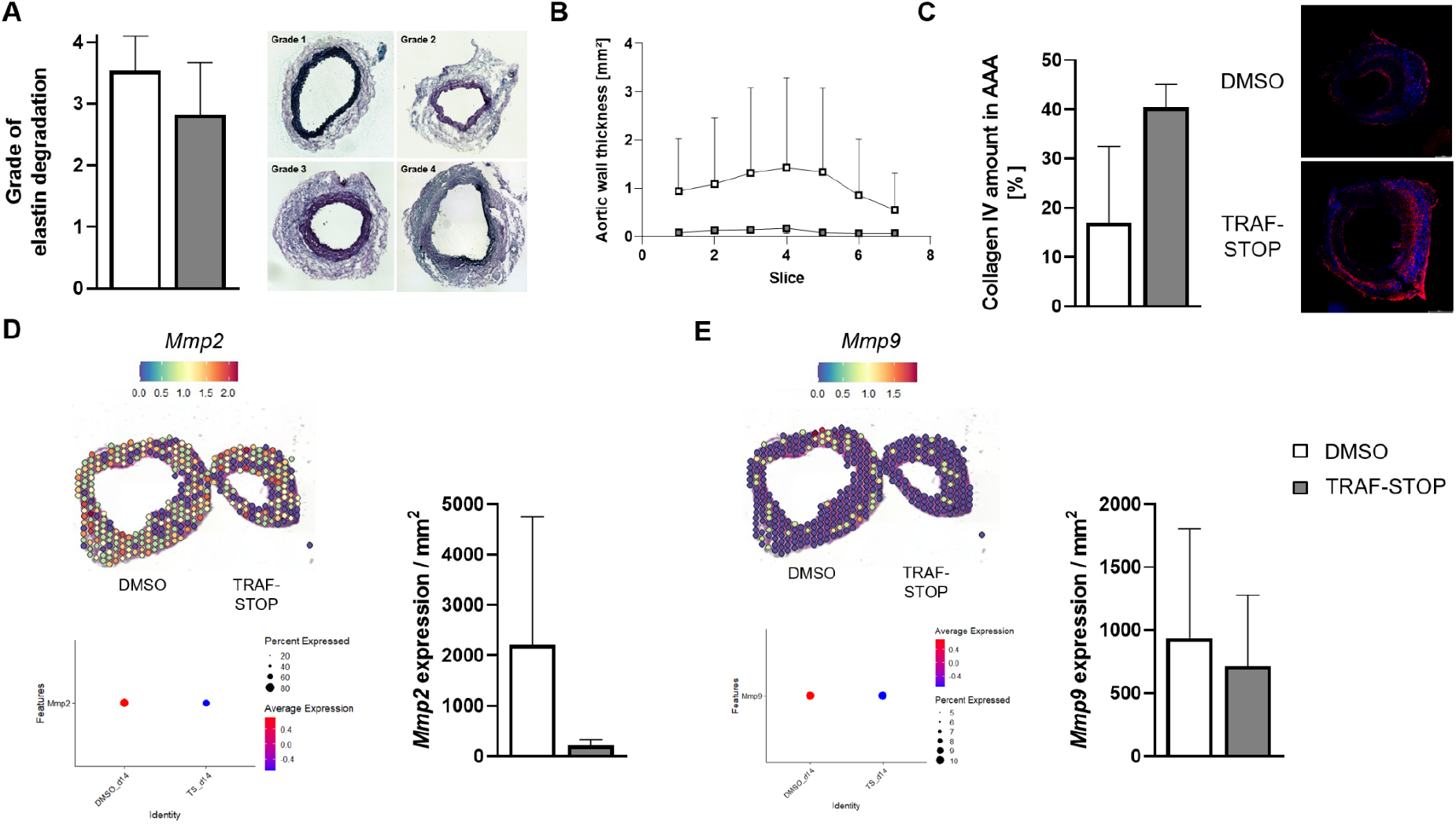
Inhibition of CD40 signaling by TRAF-STOP protects from aortic structural remodeling by a reduction of MMP2 and MMP9 expression and preserved collagen degradation. Based on an Elastica van Gieson staining, elastin degradation and aortic wall thickness were determined in 7 sequential AAA cross-sections from DMSO and TRAF-STOP-treated mice 28 days after AAA induction. (A) Quantification of medial elastin degradation (left panel) based on a grading system as indicated in the right panel. (B) Analysis of aortic wall thickness. The aortic wall was defined as area within AAA containing elastic fibers. Elastin degradation and aortic wall thickness was reduced upon TRAF-STOP treatment. DMSO n=4, TRAF-STOP n=4. (C) Increased collagen amount in aneurysmal tissue in TRAF-STOP-treated mice 28 days after PPE-induced AAA formation (left panel) and representative collagen type IV staining in AAA sections (right panel). Scale bar = 200µm; DMSO n=2, TRAF-STOP n=2. (D) Analysis of *Mmp2* gene expression by Spatial Transcriptomics (left panel) and RNAscope™ technology (right panel) 14 days after AAA induction. (E) Analysis of *Mmp9* gene expression by Spatial Transcriptomics (left panel) and RNAscope™ technology (right panel) in aneurysmal tissue 14 days post PPE-induced AAA formation. Spatial Transcriptomics: DMSO n=1, TRAF-STOP n=1. RNAscope™: DMSO n=2, TRAF-STOP n=2.

### Inhibition of CD40 signaling alters proportion of CD40^+^ macrophages and reduces TNFα expression

Analysis by Cellular Indexing of Transcriptomes and Epitopes by sequencing (CITESeq) identified a subpopulation of macrophages that highly express *Cd40* in aneurysmal tissue 7 days after AAA induction. We observed a reduction in the proportion of CD40^+^ macrophages (CD40^+^ Mac) in TRAF-STOP-treated mice compared to solvent control 7 days post PPE-induced AAA formation (**Figure 3A**). A detailed gene expression analysis revealed an upregulation of the gene C-type lectin domain family 4, member a3 (*Clec4a3*) and a downregulation of the gene Interferon induced protein with tetratricopeptide repeats 2 (*Ifit2*) within the CD40^+^ macrophages, in TRAF-STOP treated mice 7 days post PPE surgery (**Figure 3B**). We could confirm this finding, by performing Spatial Transcriptomics experiments of AAA cross-sections. TRAF-STOP treated mice showed an increased expression of *Clec4a3* within the aneurysmal tissue 7 days post AAA induction (**Figure 3C**). As *Clec4a3* signaling is involved in TNF? suppression, we investigated *Tnfa* gene expression by spatial transcriptomics and RNAscope™ (**Figure 3D**). Indeed, TRAF-STOP treated mice showed reduced gene expression of *Tnfa* in aneurysmal tissue 7 and 14 days post PPE-induced AAA formation.

**Figure 3.**
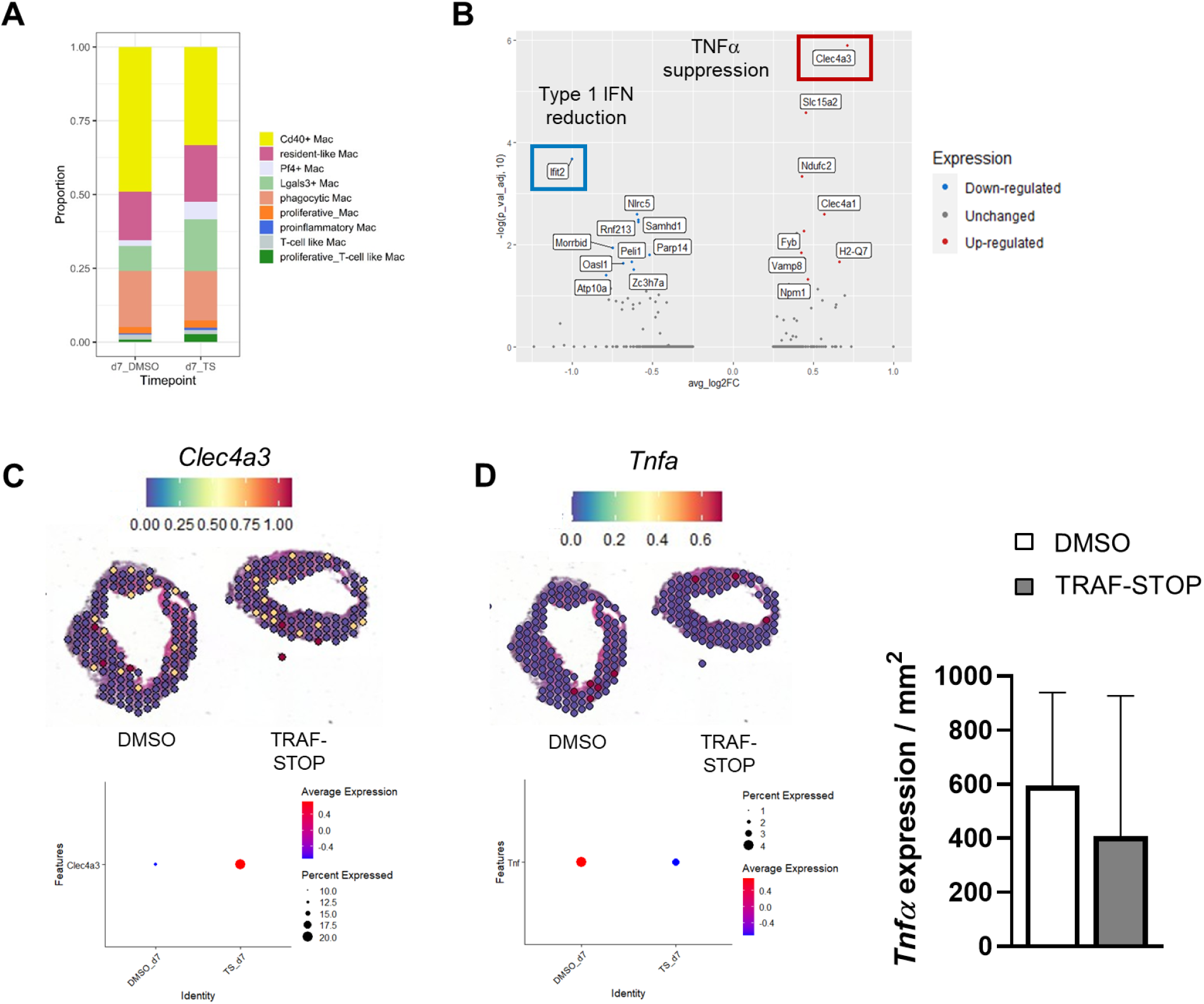
Inhibition of CD40 signaling alters proportion of CD40^+^ macrophages and reduces TNFα expression. (A) Proportion of macrophage subpopulations in aneurysmal tissue detected by CITE Sequencing of isolated cells from infrarenal abdominal aortae from C57BL/6 mice 7 days after AAA induction. TRAF-STOP treatment reduces the proportion of CD40^+^ macrophages compared to solvent control. (B) Differentially expressed genes of CD40^+^ macrophages by comparison of DMSO vs TRAF-STOP treatment, blue = down-regulated genes, red = up-regulated genes. DMSO n=1 from 2 pooled aortae, TRAF-STOP n=1 from 2 pooled aortae. Analysis of (C) *Clec4a3* gene expression by Spatial Transcriptomics and (D) *Tnfα* gene expression by Spatial Transcriptomics (left panel) in aneurysmal tissue 7 days post PPE-induced AAA formation and RNAscope™ technology (right panel) in AAA 14 days after AAA induction. Spatial Transcriptomics: DMSO n=1, TRAF-STOP n=1; RNAscope™: DMSO n=2, TRAF-STOP n=2.

## 4. Discussion

This study demonstrates that the pharmacological inhibition of CD40 signaling via its adaptor molecule TRAF-6, that is predominantly utilized in monocytes and macrophages, by the small molecule inhibitor TRAF-STOP protects from AAA progression. We could show that administration of TRAF-STOP into C57BL/6J mice reduces AAA diameter, size and the incidence of AAA development after PPE surgery. TRAF-STOP treatment stabilizes aortic wall architecture, particularly in the early phase of AAA development between day 7 and day 14 post AAA induction. The aortic wall structure is preserved by halted elastin and collagen type IV degradation, arising from a reduction of MMP2 and MMP9 expression. Accompanying, CITESeq analysis identified a CD40^+^ subpopulation of macrophages that is reduced in TRAF-STOP treated mice and shows an altered gene signature compared to mice treated with solvent control. In particular, inhibition of the CD40 signaling pathway leads to an upregulation of *Clec4a3* and reduced pro-inflammatory *Tnfα* expression in aneurysmal tissue. In addition, a downregulation of *Ifit2* expression was observed within the CD40^+^ macrophage subset in aneurysmal tissue of mice treated with TRAF-STOP.

MMPs like MMP2 and MMP9 are critically involved during AAA development leading to extracellular matrix and elastin degradation which results in aortic dilation.^18^ Moreover, AAA rupture is highly associated with increased MMP9 activity.^19^ In this study, we showed reduced elastic fiber degradation and less MMP2 and MMP9 expression in aneurysmal tissue upon TRAF-STOP treatment. Within our previous study we could show that CD40L deficient mice were protected against dissecting aneurysm formation and showed a decrease in the incidence of fatal rupture due to a decreased accumulation and activation of inflammatory cells and a reduced MMP2 and MMP9 activity in the arterial wall.^15^ This is in line with the study of Nagashima et al. that shows a reduced MMP2 mRNA and protein expression a in cultured human AAA tissue after inhibition of the CD40-CD40L pathway by trapidil.^20^

MMP2 and MMP9, both members of the gelatinase family of proteases, are known to degrade elastin as well as several collagen types including collagen type I and type IV.^21^ Accordingly, we investigated collagen type IV content in aneurysmal tissue. TRAF-STOP treatment halted collagen type IV degradation compared to solvent control. Interestingly, a recent study has demonstrated that collagen type IV deficiency promotes PPE-induced AAA formation and that the level of collagen type IV in arteries is a critical determinant for AAA formation.^22^ This is in line with our findings, implying that inhibition of CD40 signaling reduces MMP2 and MMP9 expression, which subsequently prevents the degradation of collagen type IV. Collagen plays a crucial role in repairing and regenerating the vessel wall, and also contributes to its structural integrity and strength while limiting distension at high pressure.^23^ The majority (80-90%) of collagen found in the aorta consists of fibrillar collagens type I and type III, while the remaining fraction of collagen is made up of collagens type IV, V, VI, and VII. Studies show that aneurysmal aortae in humans exhibit higher levels of collagen type I/III and collagen cross-linking, which can make arteries stiffer and more prone to dissection and rupture. On the other hand, decreased collagen content and cross-linking can weaken the aortic wall, leading to aneurysm formation and/or aortic dissection.^24^ These differences in collagen content may reflect different stages of aortic remodeling, with fibrosis occurring during later phases of inflammation in vessel repair. It is crucial to maintain a balance in collagen content for optimal aortic structure and function. In conclusion, collagen type IV seems critical in AAA development. However, as the ratio of different collagen types was shown to be important during AAA formation, other collagen types, mainly type I and type III, should be investigated in respect of a potential effect upon TRAF-STOP treatment in the future.

Different types of macrophages have been identified in aneurysmal tissue, and they play a crucial role in both the inflammatory response and the remodeling of the extracellular matrix by secreting cytokines such as TNFα, as well as MMPs.^25^ Our data identified a CD40^+^ macrophage population that shows a reduced proportion and upregulation of *Clec4a3* in aneurysmal tissue upon TRAF-STOP treatment. *Clec4a3* is predictably involved in the negative regulation of TNF production. In addition, *Tnfα* expression was reduced in aneurysmal tissue upon TRAF-STOP treatment indicating that CD40 signaling might modulate macrophages within aneurysmal tissue towards an anti-inflammatory phenotype. A previous study demonstrated that TNFα plays a central role in regulating matrix remodeling and inflammation in the aortic wall leading to AAA. In particular, Infliximab, a TNFα antagonist, decreased elastic fiber disruption, reduced macrophage infiltration, and limited the expression of MMP-2 and MMP-9 in murine aortic tissue.^26^ Furthermore, we found a downregulation of *Ifit2* within the CD40^+^ macrophage population in aneurysmal tissue of TRAF-STOP-treated mice. It has already been shown that IFIT2 plays a crucial role in the production of pro-inflammatory cytokines, as its deficiency in bone marrow-derived macrophages reduces TNFα and IL-6 secretion.^27^ In contrast, *IFIT2* gene expression was downregulated in human aneurysmal tissue from patients with stable AAA compared to patients with ruptured AAA.^28^ However, the exact role of IFIT2 in the formation and progression of AAA is poorly understood and needs to be addressed in future studies.

In summary, TRAF-STOP-treated mice develop smaller, less severe AAA and have a protected aortic architecture, reduced elastic fiber fragmentation, reduced expression of MMP2 and MMP9, and a preserved collagen IV content. This is accompanied by the reduction of pro-inflammatory processes, as *Tnfα* expression was reduced in aneurysmal tissues upon TRAF-STOP treatment. Modulation of the inflammatory and matrix-degrading phenotypes of macrophages by TRAF-STOP is a promising approach for future clinical applications.

## 5. Acknowledgements

Computational infrastructure and support were provided by the Centre for Information and Media Technology at Heinrich Heine University Düsseldorf. We thank Tobias Lautwein, Karl Köhrer, Katharina Bottermann, Dan Gorski and Jens Fischer for their help during the performance of transcriptomics experiments. We would also like to acknowledge the assistance of Katarina Raba at the Core Flow Cytometry Facility at the Institute for Transplantation Diagnostics and Cell Therapeutics Düsseldorf.

